# Levels of Representation in a Deep Learning Model of Categorization

**DOI:** 10.1101/626374

**Authors:** Olivia Guest, Bradley C. Love

**Affiliations:** Department of Experimental Psychology, University College London; Department of Experimental Psychology, University College London, and The Alan Turing Institute

**Keywords:** categorization model, deep networks, pigeons, ventral stream, representational hierarchy

## Abstract

Deep convolutional neural networks (DCNNs) rival humans in object recognition. The layers (or levels of representation) in DCNNs have been successfully aligned with processing stages along the ventral stream for visual processing. Here, we propose a model of concept learning that uses visual representations from these networks to build memory representations of novel categories, which may rely on the medial temporal lobe (MTL) and medial prefrontal cortex (mPFC). Our approach opens up two possibilities: *a*) formal investigations can involve photographic stimuli as opposed to stimuli handcrafted and coded by the experimenter; *b*) model comparison can determine which level of representation within a DCNN a learner is using during categorization decisions. Pursuing the latter point, DCNNs suggest that the shape bias in children relies on representations at more advanced network layers whereas a learner that relied on lower network layers would display a color bias. These results confirm the role of natural statistics in the shape bias (i.e., shape is predictive of category membership) while highlighting that the type of statistics matter, i.e., those from lower or higher levels of representation. We use the same approach to provide evidence that pigeons performing seemingly sophisticated categorization of complex imagery may in fact be relying on representations that are very low-level (i.e., retinotopic). Although complex features, such as shape, relatively predominate at more advanced network layers, even simple features, such as spatial frequency and orientation, are better represented at the more advanced layers, contrary to a standard hierarchical view.

---

Deep convolutional neural networks (DCNNs) excel in image classification classifiers tasks, having set a new standard in machine learning benchmarks Krizhevsky, Sutskever, and Hinton (2012). These models are trained to predict category labels (e.g., “crow”, “sunset”, “car”) from visual inputs (e.g., photographs) through supervised learning. After training, DCNNs can generalize to unseen images from a test set (Russakovsky et al., 2015). DCNNs rival human performance when tested on specific object recognition tasks (Krizhevsky et al., 2012; LeCun, Bengio, & Hinton, 2015). These large networks, with tens of layers and thousands of connection weights, build on a tradition of connectionist modeling in Psychology that dates back to pioneers such as Rosenblatt (1958), J. A. Anderson (1972), and Rumelhart, Hinton, and Williams (1986).

Unlike the fully connected networks that were popular in 1980s, convolutional networks are based on general organizational principles of the visual cortex (Fukushima & Miyake, 1982; Lecun & Bengio, 1995; LeCun et al., 1990; Mozer, 1991). DCNNs models learn filters (i.e., receptive fields for certain features) that are applied to a portion of that network layer, corresponding to a location in the stimulus image. For example, at an early network layer, a learned filter may be somewhat analogous to an orientation column found in the visual cortex. At more advanced layers, the network’s receptive fields become more global, covering a greater extent of the image representation. The features at these more advanced layers can be characterized as being more abstract, complex, and invariant to lower-level image properties.

One hypothesis is that the layers in DCNNs bear some relationship to different regions along the ventral stream. In support of this hypothesis, DCNNs offer the best account of neural activity along the ventral stream with the progression of model layers roughly corresponding to processing stages along the ventral stream (Cichy, Khosla, Pantazis, Torralba, & Oliva, 2016; Güçlü & van Gerven, 2015; Khaligh-Razavi & Kriegeskorte, 2014; Schrimpf et al., 2018; Yamins & DiCarlo, 2016; Yamins et al., 2014). Beyond accounting for neural measures, DCNNs have also proven to be valuable theoretically in terms of understanding human behavior (Kubilius, Bracci, & Op de Beeck, 2016; Lake, Zaremba, Fergus, & Gureckis, 2015; Ritter, Barrett, Santoro, & Botvinick, 2017). Thus, DCNNs are a potential bridge that can link levels of representation in the brain to behavior (Love, 2016).

In this paper, we present a new model of category learning that uses representations of visual stimuli from a DCNN layer. By using representations from a DCNN, we are not restricted to the hand-coded representations typically used in psychological models of category learning (e.g., J. R. Anderson, 1990; Love, Medin, & Gureckis, 2004; Nosofsky, 1986) and can instead apply the model to naturalistic stimuli (e.g., Nosofsky, Sanders, Gerdom, Douglas, & McDaniel, 2017). By comparing model fits that operate on representations from different DCNN layers (e.g., category learning models using early vs. late DCNN layers), we can infer which types of representations a learner used.

Previous work in the cognitive neuroscience of category learning has relied on stimulus representations specified by the experimenter (Davis, Love, & Preston, 2011, 2012). Using these stimulus representations, an array of cognitive models have been evaluated as high-level accounts of how the brain acquires concepts, including the exemplar model used here (Mack, Love, & Preston, 2016; Mack, Preston, & Love, 2013). One basic hypothesis is that learning novel categories depends on structures in the Medial Temporal Lobe (MTL), including the hippocampus, that work in concert with medial prefrontal cortex (mPFC) to identify category-relevant information (Love & Gureckis, 2007; Mack et al., 2016; Mack, Love, & Preston, 2018; Mack, Preston, & Love, 2017). Although this characterization is human centric, there may be general categorization processes shared across species (Smith, Minda, & Washburn, 2004; Soto & Wasserman, 2012).

In some senses, research in category learning focuses of learning mechanisms that sit atop of the ventral stream and, therefore, can be viewed as complementary to work in deep learning that has been related to the ventral stream. Category learning models aim to capture how people and animals rapidly acquire novel concepts based on small set of examples. To accomplish this feat, they require useful stimulus representations of the visual input. Setting the final layer of a DCNNs aside, which is not used in the simulations reported here, a DCNN’s job in object recognition ends where a category learnng model’s job begins. In terms of implementation, these two models classes are hypothesized to rely on non-overlapping brain regions for the most part.

Thus, it’s natural from a neural computational perspectve to combine the strengths of these two models to characterize concept learning with real-world stimuli. The strength of the DCNNs are in building visual representations at multiple levels (or layers) from *prior* training on hundred of thousands of ecologically-valid stimuli that approximate the natural image statistics from a lifetime of visual experience. The strength of category learning models is capturing how agents rapidly detect and store patterns of experiences to acquire novel concepts. DCNNs can characterize the inputs to category learning models, allowing for an end-to-end account of categorization behavior and its neural underpinnings.

We do not discount the possibility of lifelong tuning of visual representations, but instead assume that the ventral stream does not radically change over the course of a brief categorization learning study. Thus, for the simulations reported here, we treat the DCNN model as fixed. All learning takes place within the category learning model which take visual representations from the DCNN as input. For example, in learning two classify sparrows and robins, the exemplar model would store away experiences in its memory using visual representations provided by the DCNN. The DCNN provides the coding of these stimuli, which spares the experimenter from hand-coding stimuli in terms of features (e.g., “red”, “has wings”, “small”, etc.).

Likewise, we do not discount that visual representations in the ventral stream can be modulated by top-down influences during category learning. Indeed, relevant aspects of visual representations are accentuated in the ventral stream during category learning (Bobadilla-Suarez, Ahlheim, Mehrotra, Panos, & Love, 2019; Braunlich & Love, 2018; Folstein, Palmeri, Van Gulick, & Gauthier, 2015).

At this early juncture in this research program, we do not address these non-feedforward processes. As will become clearer, one advantage of setting these important questions aside for the moment is that we can better characterize the representational biases present across DCNN layers absent top-down task modulation.

In summary, we use DCNNs as models of the ventral stream that provide visual representations that we take to reflect a lifetime of visual experience. These visual representations serve as inputs to an exemplar model of categorization (proposed to rely on the MTL and mPFC) that can rapidly learn novel concepts over a handful of trials. As discussed below, our work allows us to characterize the nature of visual representations at various layers of DCNNs and evaluate how they contribute to novel concept learning through fits of the exemplar model. Below, is an overview of the simulations.

First, we adopted this approach to understand children’s shape bias, that is their tendency to generalize according to shape as opposed to other stimulus dimensions such as color (L. B. Cohen & Cashon, 2003; Gershkoff-Stowe & Smith, 2004; Landau, Smith, & Jones, 1992, 1988; Quinn & Eimas, 1996; Rakison & Butterworth, 1998; Samuelson, Horst, Schutte, & Dobbertin, 2008; Samuelson & Smith, 1999, 2000). Working with naturalistic stimuli, we found that the shape bias is consistent with use of visual representations from more advanced network layers. In this fashion, rather than simply conclude that the shape bias may in part be attributable to the statistics of visual experience, we can instead make inferences about which statistics are key (e.g., statistics that involve higher-level visual features).

Subsequently, we conducted simulations with a well-controlled stimulus set to better understand how various types of features are encoded at different network layers. We concluded that, while abstract shape features are relatively more important at advanced network layers, all features, even basic features like orientation and spatial frequency, are best encoded at later network layers.

Our finding that a range of features are best encoded at later network layers contrasts with classical hierarchical views of visual object representation, which assume that feature encoding follows a strict ordering with low-level features preceding higher-levels features without both feature types coexisting at the same representational layer. For example, according to a strict hierarchical view, simple visual features, like lines and colors, are combined to form increasingly more complex features (Serre & Poggio, 2010).

However, our findings are in agreement with findings from visual neuroscience. Although there is evidence that low-level visual properties, such as orientation, frequency, and position are encoded in early visual areas (Hubel, 1963; Hubel & Wiesel, 1962; Watson, Hartley, & Andrews, 2014) and features invariant to differences in these lower-level features are encoded in latter downstream areas (Bracci & Op de Beeck, 2016; M. A. Cohen, Alvarez, Nakayama, & Konkle, 2016; Grill-Spector et al., 1999; Haushofer, Livingstone, & Kanwisher, 2008; Op de Beeck, Torfs, & Wagemans, 2008), these findings alone are not uniquely supportive of a strict hierarchical account. Indeed, these findings are consistent with our simulations, as are recent finding demonstrating that even lower-level features are encoded, in some cases better, in latter brain regions (Ahlheim & Love, 2018; Hong, Yamins, Majaj, & DiCarlo, 2016; Lescroart & Gallant, 2019; Lu et al., 2018; Rice, Watson, Hartley, & Andrews, 2014). These findings and our results suggest that functionally there is an inverted pyramid in which more features are encoded at more advanced network (and brain) layers.

Finally, with this better understanding of how DCNN encode visual features, we applied our model to a visual categorization study involving pigeons. The pigeons were trained to classify medical images and, impressively, they reach performance levels matching those of human expert technicians (Love, Guest, Slomka, Navarro, & Wasserman, 2017). However, our modeling demonstrates that the pigeons may be relying on representations that are very low-level (i.e., retinotopic statistics) to base their decision-making. These results highlight how our categorization model using DCNN visual representations could be used to understand the information (statistics) that a learner relies upon. We will characterize what is represented at various levels in the DCNN, and then use our model to understand the level of representation used in a given categorization task. Before presenting these results, we first present the formalism for the DCNN and our categorization model which makes use of these representations.

## Modeling Categorization using DCNN Representations

We model categorization using an exemplar approach with exemplar representations coming from a pre-trained DCNN (see Figure 1), as opposed to relying on hand-coded representations from the experimenter. In this section, we formalize this approach, which we later use to understand shape bias and characterize how pigeons classify complex visual stimuli.

**Figure 1.**
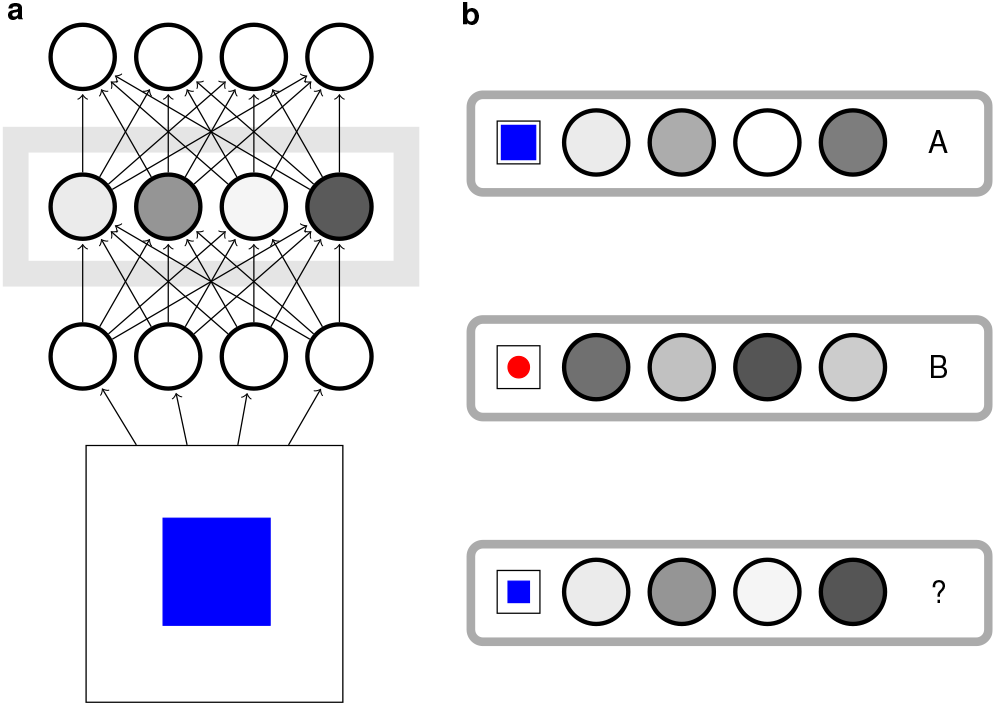
A schematic of the DCNN-plus-exemplar model. On the left in **a**), a stimulus is presented to the pre-trained DCNN, which leads to pattern of activity at the network layer of interest. This pattern provides the exemplar representation for that stimulus. On the right in **b**), the exemplar model has two previously stored exemplars, one each from category “A” and “B”. For a typical learning task, there would be many more exemplars stored (one for each training trial). The similarity between the current stimulus and all stored exemplar representations determines how the current stimulus is classified.

### DCNN Details

The DCNN is used as a way to extract features from realistic stimulus representations, e.g., photographs. The convolutional network used herein is Inception-v3 GoogLeNet, a DCNN classifier (Szegedy, Liu, et al., 2015; Szegedy, Vanhoucke, Ioffe, Shlens, & Wojna, 2015). The output layer is trained to represent very high-level conceptual categories (1000 mutually exclusive classes, e.g., sunglasses, moped, jellyfish, etc., Russakovsky et al., 2015). Although these output classes do not contain options for the specific stimuli we use, the network provides a distributed answer across these 1000 units/categories thus being able to solve a classification task of unknown categories. In the following sections, we will use the activations of each layer (e.g., the output layer, as well as all layers below it) to embed each input stimulus. Each embedding space has a similarity structure matching the distinction in the inputs, i.e., functional smoothness holds, for the deeper layers (e.g., roughly two thirds depth and onwards; Guest & Love, 2017). Thus, if items are similar in behavioral ratings space, e.g., a lion and a tiger, their representations will be similar in the appropriate layers’ embedding spaces.

### Categorization Model

The pattern of activity elicited by a stimulus in the DCNN layer of interest provides the exemplar representation of a stimulus. A standard exemplar model of categorization (cf., Medin & Schaffer, 1978; Nosofsky, 1986, utilizes these representations).

Formally, the exemplar model determines the probability that stimulus *S*_*i*_ is in category *j* by calculating the similarity sim(*S*_*i*_, *S*_*j*_) to exemplars from *j* and those of the contrasting categories,

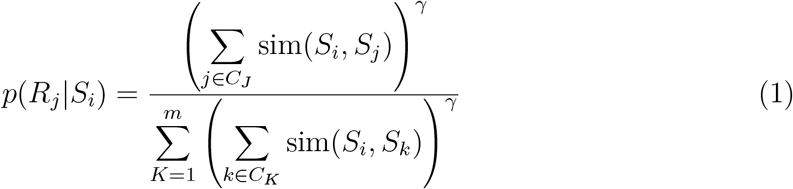

where *C*_*J*_ denotes category *J*, *m* is the number of categories the model has learned, and *γ* is a decision parameter (based on: Maddox & Ashby, 1993; Nosofsky, 1986; Nosofsky & Zaki, 2002). When *γ* is set to values greater than 1 the model computes a max response (e.g., yes or no) and when it is set to 1 more fine grained responses will be given (e.g., a probability). The decision parameter *γ* can be fit to the data as done in previous work using the classic generalized context model (GCM; Maddox & Ashby, 1993; Nosofsky & Zaki, 2002). To avoid negative similarities (Guest & Love, 2017), Pearson correlation plus 1 is used as the similarity function,

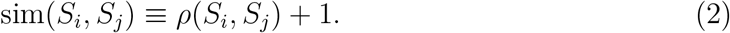

We leave consideration of attention-weighted similarity measures for future work (cf. Lindsay & Miller, 2018; Nosofsky & Zaki, 2002).

## Shape Bias Simulations

The shape bias refers to children’s tendency under certain circumstances to generalize according to shape (L. B. Cohen & Cashon, 2003; Gershkoff-Stowe & Smith, 2004; Landau et al., 1992, 1988; Quinn & Eimas, 1996; Rakison & Butterworth, 1998; Samuelson et al., 2008; Samuelson & Smith, 1999, 2000). We assessed shape bias using a triplet task (see Figure 2a) in which participants, or models in our case, are asked to choose whether the shape or color match stimulus is more similar to the probe stimulus.

**Figure 2.**
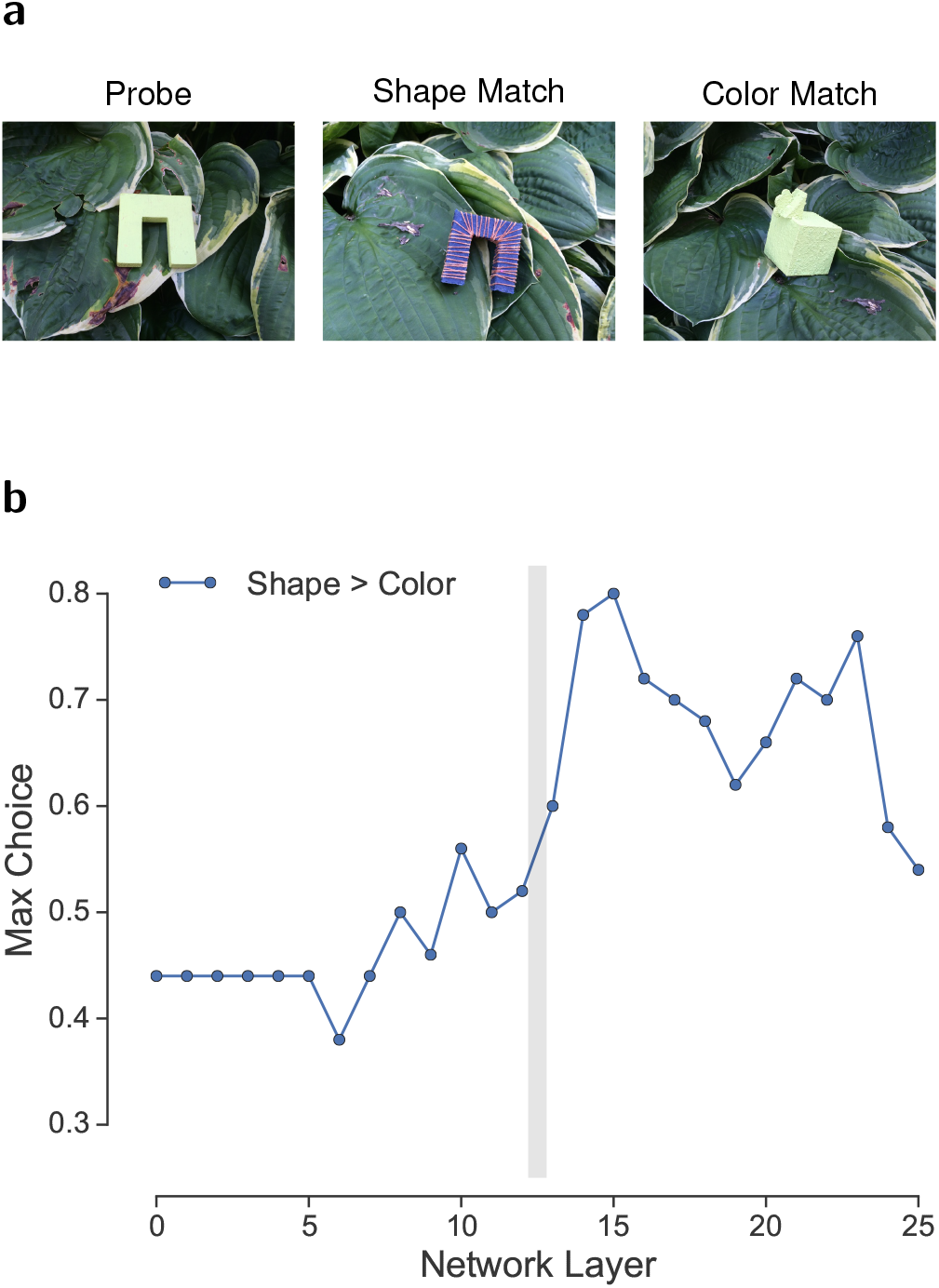
The stimuli used and the max choice accuracy in the triplet task. **a**) Stimuli used in the triplet task. On the leftmost column is the probe or standard item, while to its right are the objects that share the same shape (middle column, shape match) and color (right column, color match). **b**) Preference for shape over color match in the triplet task for each layer of the network. At the early layers, to the left of the vertical gray line there is a significant preference for the stimulus that matches the probe in color. At the later layers, to the right of the vertical gray line, the network prefers to categorize using shape.

We used this triplet task to assess shape (or color) bias in DCNNs, which in turn we can use to infer the types of statistics that humans rely upon when they exhibit related biases. Our simulations conceptually replicate and extend those by Ritter et al. (2017) which focused on the outputs of DCNNS. Our key extension is to consider all DCNN layers such that we can characterize the entire range of network representations from pixel-based (akin to the retina) to higher-level representations (akin to inferotemporal cortex).

Thus, the task was modeled according to the preceding equations, but with different simulations relying on representations from different DCNN layers for computing similarity (i.e., Equation 2). The two match stimuli, see Figure 2a, were considered training examples of contrasting categories, whereas the probe stimulus was used as the test stimulus. To simplify analysis, the max choice was taken as the model’s response, which corresponds to setting the *γ* response parameter to positive infinity.

### Stimulus Set

The stimuli consist of a set of 50 triplets, 10 unique objects photographed with 5 different backgrounds by Linda Smith’s lab (referred to as the CogPsyc dataset in Ritter et al., 2017). An example triplet with the “green leaf” background is shown in Figure 2a. The set is available to download at http://www.indiana.edu/~cogdev/SB_testsets.html (as document files, containing a number of images per page) as well as in the OSF repository: https://osf.io/jxavn/ (as individual image files).

### Results and Discussion

For early network layers, the probe stimulus was similar to the color match than the shape match, whereas at layer 13 this preference reversed with the network displaying a shape bias (see Figure 2b). There is a clear transition in bias, delineated by the gray band shown in Figure 2b, between the early versus the late layers which is statistically reliable, *χ*^2^(1, *N* = 1300) = 63.34, *p* < 0.0001. These results indicate that shape bias is likely driven by higher-level statistics and that reliance on lower-level representations can lead to other biases, such as a color bias.

## Systematic Study of Deep Learning Representations

In the previous simulations, we found that shape matches are more consequential than color matches using representations from more advanced network layers. In this series of simulations, we use well-controlled stimuli to systematically evaluate the coding and relative influence of various types of features, such as shape, size, and color, across network layers. We also consider how positionally independent the network’s representations are. The purpose of these simulations is to better characterize the representations at various network layers.

### Stimulus Set

#### Overlapping Stimuli and Category Structure

The stimuli, shown in Figure 3, have three binary-valued dimensions: shape (square or circle), color (red or blue), and size (big or small). The stimulus set is referred to as overlapping because each stimulus is spatially centered in the middle of the image. We also consider an alternative stimulus set in which the color dimension is replaced by monochrome light gray or dark grey stimuli. This stimulus set is akin to those used in studies of human category learning. However, whereas each stimulus would be coded by experimenters and modelers as a vector with three binary-valued dimensions, here we work with directly with the uninterpreted pixel-values as shown in Figure 3.

**Figure 3.**
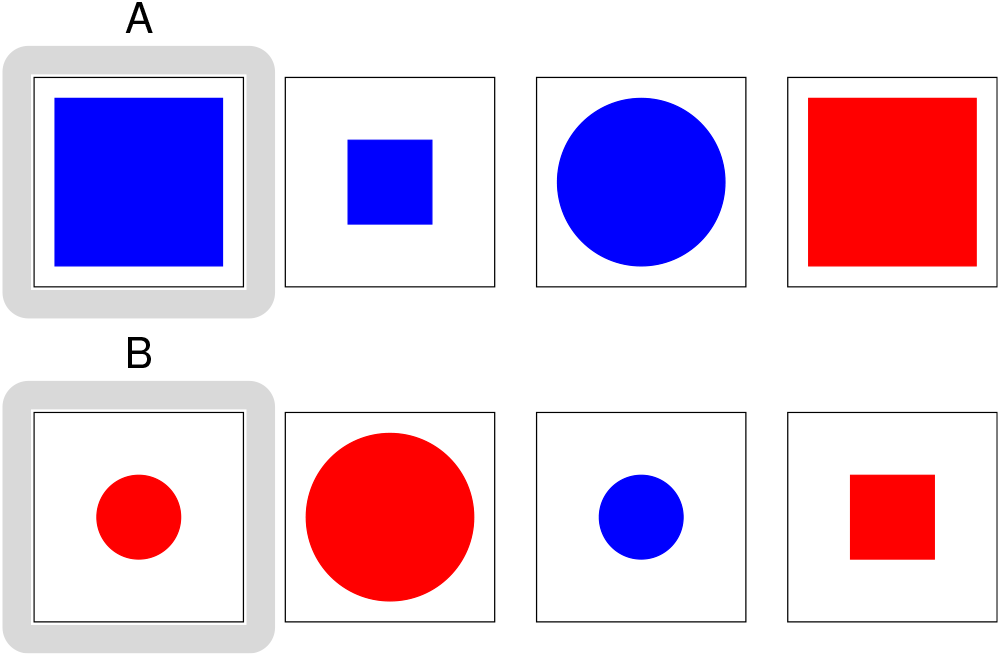
The 3-dimensional binary-valued stimulus set. Stimuli are members of the category for which they match the prototype on 2 or more of the three features (size, shape, and color). The prototype for each category is shown with a gray outline in the first column.

Each category is centered around a prototype and the two category prototypes display contrasting values on each dimension. In the case where, for example, the big blue square is assigned as the “prototype” for category A (denoted by a gray outline in Figure 3) then the members of A will be the items which share 2 out of 3 dimension values with the prototype, shown on the top row of Figure 3. By the same token, the members of category B are defined, shown on the bottom row of Figure 3. Given we defined A’s prototype and that the two categories’ prototypes must have nothing in common, the prototype of category B can only be the small red circle. The prototype role can be played by any stimulus, resulting in 4 unique possible permutations for the two categories.

Training consists of only two training items, the two prototypes. The remaining 6 items are used as test (generalization) items from which we evaluate model performance. As a reminder, for all simulations, it is the exemplar category learning model that learns during these brief tasks using the visual representations provided by the DCNN that previously has been trained on hundreds of thousands of real-world stimuli, which we take to reflect a lifetime of visual experience.

#### Non-overlapping Stimuli and Category Structure

The non-overlapping stimuli are identical to the overlapping set with the exception that the spatial overlap between training and test items is removed. In particular, one prototype is presented on the left side of the image and the other on the right. Test items are presented in a non-overlapping position in the center of the image. These stimuli allow us to assess whether network representations are positionally invariant or are determined by low-level (pixel) overlap.

### Results and Discussion

Figure 4b shows that with non-overlapping stimuli, whether they are color or greyscale, that accuracy on test items rises as one ascends network layers. In contrast, for overlapping stimuli (see Figure 4a) there is a U-shaped pattern with performance dipping and then finally rising at more advanced network layers. For both overlapping and non-overlapping stimulus sets, there is a curious pattern in which the colored stimuli have an advantage over the grayscale stimuli even though both are binary-valued and therefore appear informationally equivalent.

**Figure 4.**
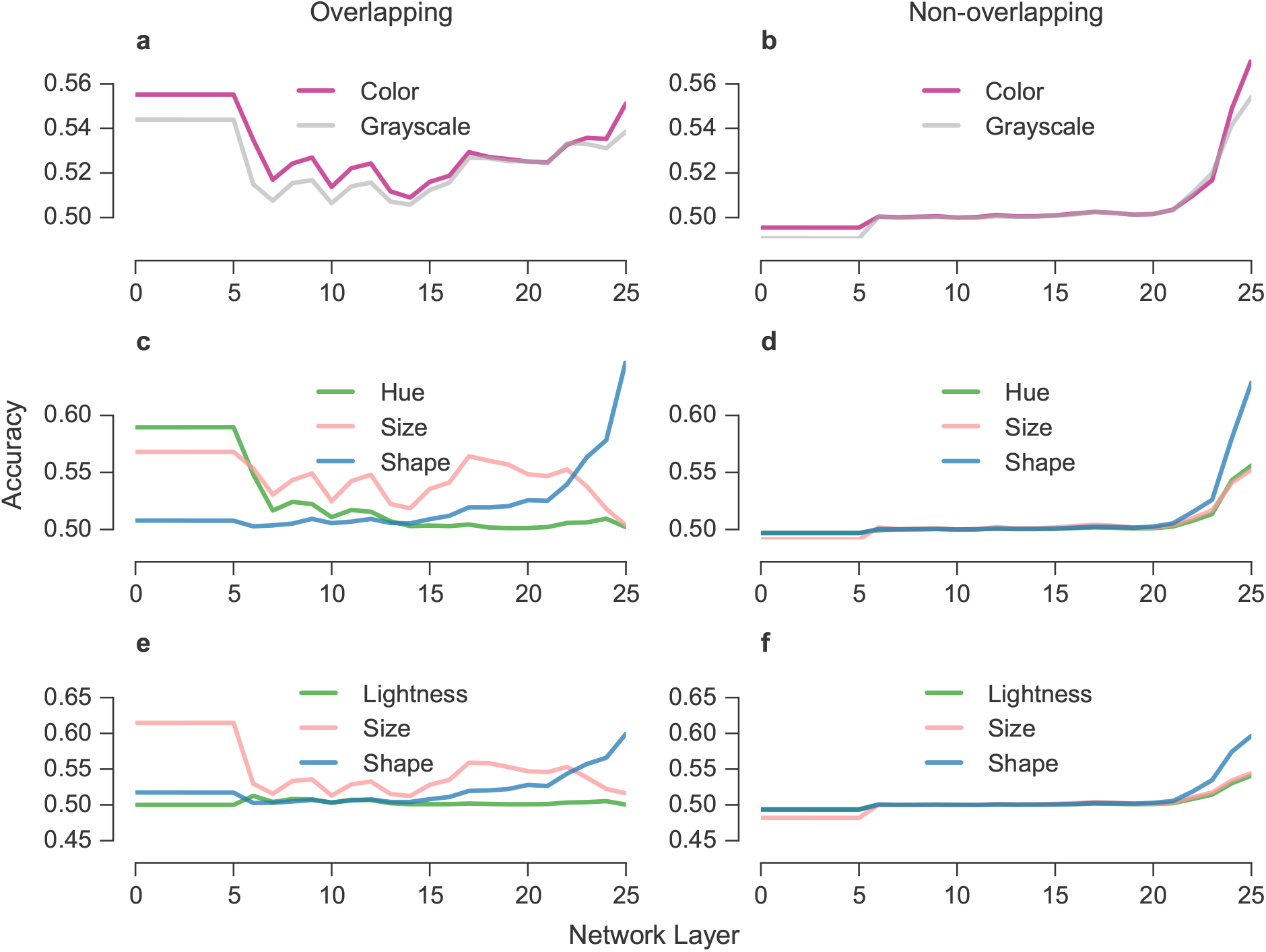
Accuracy (with the *γ* decision parameter set to 1) for the exemplar model using various network layers to provide exemplar representations. In the top row, accuracy is shown: **a**) when using the overlapping pattern set in both grayscale and color and **b**) when using the non-overlapping pattern set in both grayscale and color. In the middle row, accuracy is by dimension match to the category prototype for **c**) color overlapping stimuli, **d**) color non-overlapping, **e**) grayscale overlapping, and **f**) grayscale non-overlapping stimuli.

These basic results can be examined at a finer-grain by considering performance for individual stimuli that match their prototype along particular stimulus dimensions. For example, if shape matches were more consequential than color matches at advanced network layers, then shape-matched stimuli should display higher accuracy. All stimulus sets show a shape advantage at advanced network layers (see Figure 4). However, the overlapping stimulus sets show a shape disadvantage at early network layers. This suggests that biases for dimensions other than shape may occur for overlapping stimuli when lower-level representations are relied upon.

Overall, these results confirm those of the previous shape-bias simulations with two important additions. First, advantages for non-shape dimensions may be driven by reliance on low-level representations that are not location invariant. Second, color has a peculiar advantage over grayscale values even though there does not appear to be any informational reason for this difference.

This curious color advantage over grayscale is due to the confluence of how color is represented in images and how similarity is calculated in the exemplar model. Each pixel location is represented by a triplet of RGB values, somewhat analogous to the three different cone types in human retina. For example a blue pixel is represented as [0 0 1], whereas a gray pixel is [0.5 0.5 0.5]. In the similarity function (Equation 2), differences across vectors with this color representation are more consequential (i.e., affect similarity more) than differences in greyscale values, because the exemplar model’s similarity function relies on Pearson correlation, which is based on angle differences like vector cosine does. These model properties prove useful in a later simulation to capture advantages pigeons show in categorizing color stimuli.

## A Hierarchy of Visual Features or an Inverted Pyramid?

In the previous section, shape was found to be more consequential than other features at more advanced network layers as representations became more location invariant. One possibility is that higher-level features like shape are represented at more advanced network layers whereas more basic features are represented lower in the hierarchy. An alternative possibility is that all features are more strongly manifested at more advanced network layers in absolute terms and that certain features, like shape, only dominate in relative terms.

To help shed light on these questions, we consider two stimulus dimensions that are considered very low-level, namely orientation and spatial frequency, that can be continuously manipulated. Using Gabor patches (see Figure 5) varying across many possible values of orientation and spatial frequency, we evaluate whether these stimuli form a two dimensional representational space in each layer of the network. To foreshadow, they do and the pattern is present at all layers.

**Figure 5.**
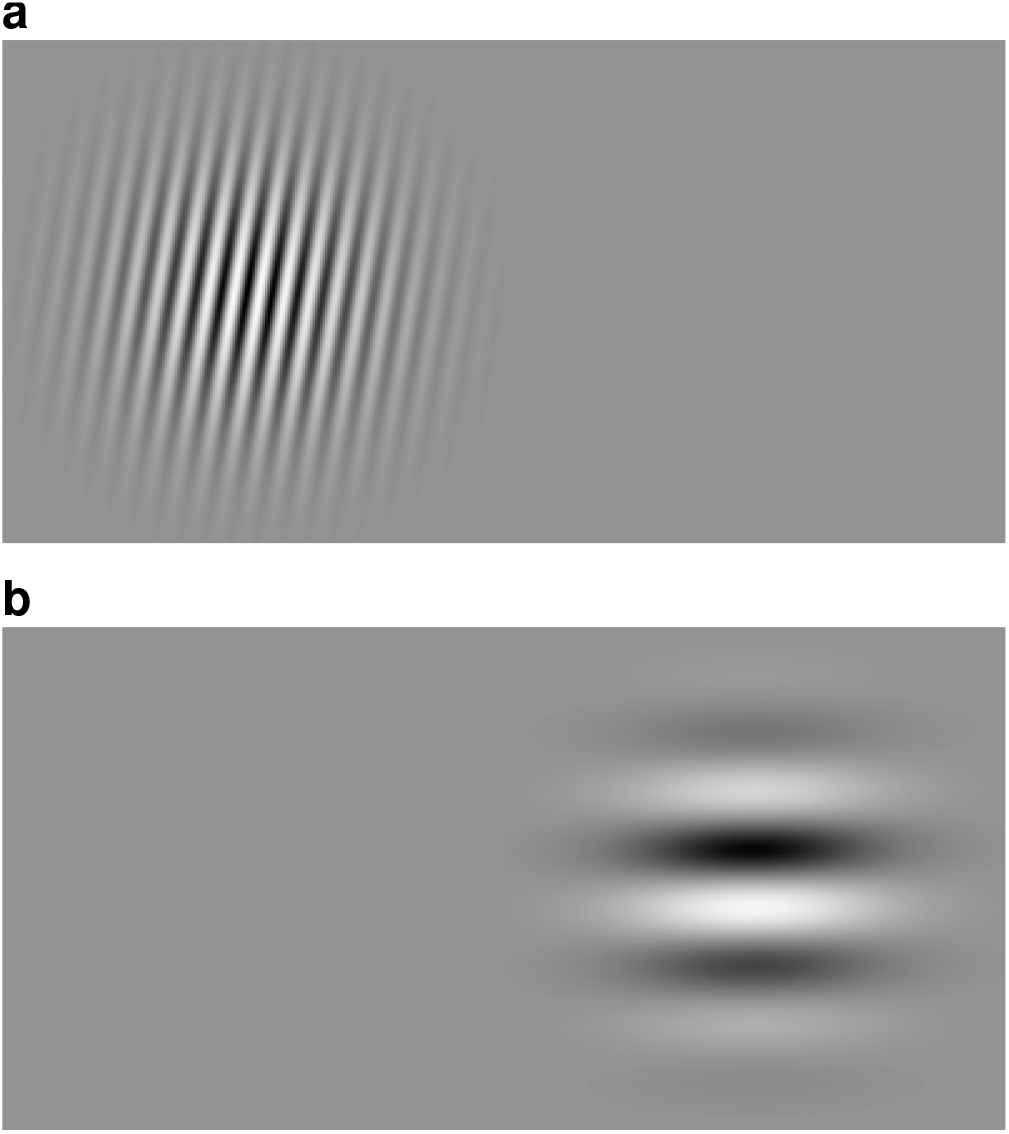
Examples of the Gabor patch stimuli that are non-overlapping. The Gabor patch on top in **a**) has a higher spatial frequency and more vertical orientation than the Gabor patch in **b**) shown below.

To evaluate at which network layer orientation and frequency are most precisely represented, we evaluate how well a Gabor patch matches a non-spatially overlapping version of itself and find that the degree of match increases across network layers. This result is inconsistent with the idea that lower-level features are represented at lower network layers. Instead, we find that precise representations of orientation and frequency are found at more advanced network layers.

### Stimulus Set

The stimuli used here are 81 Gabor patches with varying frequency and orientation. In the first simulation, Gabor patches were centered and spatially overlapped. In the second, simulation they were non-overlapping as shown in Figure 5.

### Results and Discussion

To evaluate whether each network contained a two-dimensional representational space of orientation and frequency, we performed a Principal Components Analysis (PCA) on the activation patterns of all 81 stimuli. Specifically, this PCA was done on a matrix in which each row corresponds to a stimulus and each column contains the corresponding activations of DCNN units.

We considered the first two components of the PCA solution for each layer and correlated each vector of coordinates with the experimenter determined coding of orientation angles (1–81) and frequency (10–90), resulting in 4 correlations. The two highest correlations also involved separate pairs of vectors (e.g., component 1 with frequency and component 2 with orientation). Those top 2 correlations are plotted in Figure 6.

**Figure 6.**
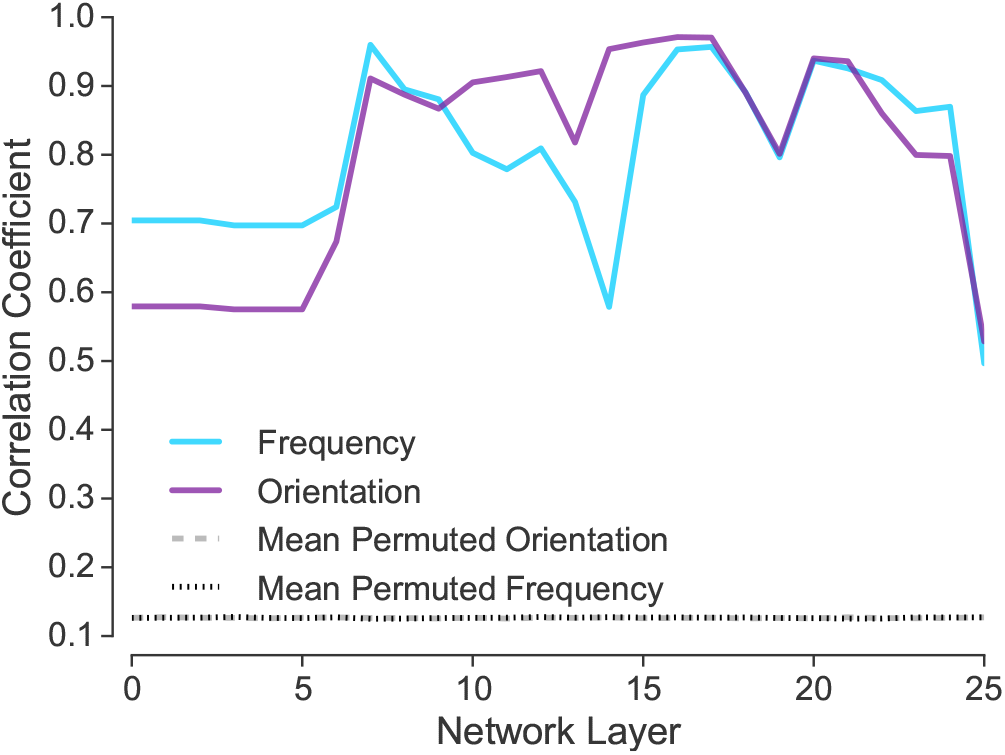
Network activity resulting from viewing Gabor patches varying in orientation and frequency was subjected to PCA. The correlation between the first two components and the frequency and orientation value of the stimulus is plotted, along with the mean of the null distribution. The results indicate that information about frequency and orientation is represented at all network layers.

At every network layer, orientation and frequency are represented. Using a permutation test, all correlations were statistically significant, *p*_*permuted*_ < 0.001. The null distribution for the permutation test was performed by repeatedly shuffling the stimulus labels and carrying out the analysis procedure.

The previous results indicate that frequency and orientation representations exist across all network layers. These features are not relegated to lower network layers. A second simulation assesses how precise orientation and frequency information is represented at each layer.

Using Gabor patches that do not spatially overlap (see Figure 5), we considered how similar a Gabor patch presented on the left side of an image was to each of the 81 possible Gabor patches presented on the right. Ideally, a Gabor patch would be most similar to the exact version of itself translated to the right. As shown in Figure 7, at more advanced network layers the left- and ride-side version of a Gabor patch tended to be most similar to each other. These results suggest that putatively lower-level features, such as orientation and frequency, are most precisely represented at more advanced network layers.

**Figure 7.**
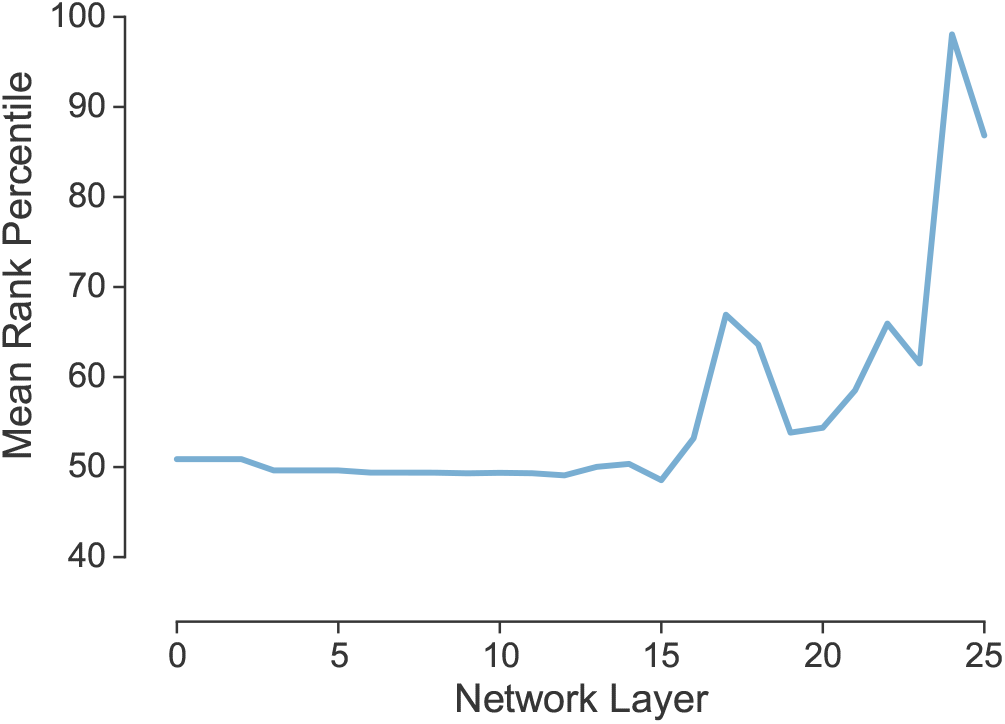
How similar identical, but non-spatially overlapping, Gabor patches are to one another relative to other stimuli. The 50*th* percentile indicates chance performance in which the matching Gabor patches are as similar to each other as would be expected for randomly selected stimuli.The 100*th* percentile indicates perfect discrimination.

## Are Pigeons Using Lower- or Higher-level Representations when Categorizing?

We have established that shape, which is a complex feature, relatively dominates in the DCNN at advanced layers, whereas more basic features, like color, relatively dominate at lower network layers. In absolute terms, all features appear to be more precisely coded at advanced network layers. We have also observed and have a plausible explanation for why color stimuli can lead to better performance than informationally-equivalent grayscale stimuli.

These findings are interesting and valuable in their own right. Additionally, understanding these basic characteristics of DCNN representations makes our categorization model (see Figure 1), which utilizes these representations, potentially more informative. In particular, we can assess whether an individual or some population is relying on lower- or higher-level visual representations with some understanding of what that would imply.

In this section, we consider whether pigeons rely on lower-level or higher-level image representations when categorizing. Pigeons are excellent at classifying visual stimuli (Bhatt, Wasserman, Reynolds, & Knauss, 1988). For example, pigeons trained to discriminate between medical images of normal and cancerous breast tissue generalized to novel stimuli and attained human-level accuracy (Levenson, Krupinski, Navarro, & Wasserman, 2015). However, it is an open question what the basis for this performance is. Are pigeons relying on very low-level pixel-like information or are they extracting and using higher-level abstract shape information? We will address this question by applying our model to a study conducted by Wasserman and colleagues. Pigeons were trained to discriminate cardiograms (see Figure 8) as normal or abnormal and then were transferred to novel stimuli during testing. These data are reported in Love et al. (2017).

**Figure 8.**
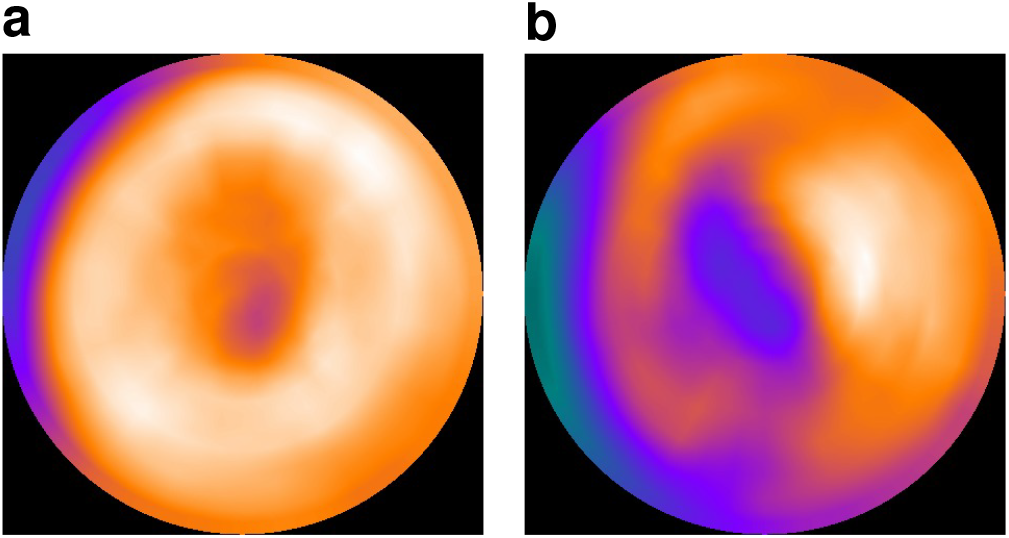
Two examples cardiogram stimuli: **a**) a normal cardiogram without any perfusion damage; and **b**) an abnormal cardiogram with total perfusion damage 20 (of a maximum of 51).

We applied our model to the same images the pigeons viewed and report how the model performs on novel test cardiogram images, qualitatively comparing the model’s performance to that of the pigeons. Pigeons were trained under three conditions: *1*) Pigeons trained on colored cardiograms and tested on colored cardiograms performed best. *2*) Pigeons trained on informationally equivalent grayscale cardiograms and tested on grayscale cardiograms performed second best. *3*) Pigeons trained on color cardiograms and tested on grayscale cardiograms performed worst. This ordering of conditions is the basic qualitative pattern of results we hope to capture with our model. Additionally, the model should, like the pigeons, show a gradient of responding to cardiograms depending on the level of perfusion damage. In particular, images with a higher-level of damage should be more likely to be classified as abnormal.

### Stimuli

The stimuli used here consist of medical images, cardiograms, in one of two classes: healthy or abnormal, see Figure 8. They have been provided by Wasserman and colleagues, who used them to train their pigeons on the same classification task as our model in this section (Love et al., 2017). There are two sets of cardiogram stimuli: colorized and grayscale. It is important to note that the output of the cardiograph is a grayscale image which is then subsequently colorised — this is a preference of the medical technicians involved in reading cardiographs. No new information is present in the colorized images as their pixels have been mapped losslessly from the original grayscale output to a colorized space.

### Results and Discussion

Our model’s performance on the three training and test conditions is shown in Figure 9 (where the optimum accurancy is shown too to control for response bias, showing what is possible working with training set). Working with DCNN representations in pixel space (the lowest-level representations, akin to retinal), our model matches the qualitative difficulty ordering that the pigeons displayed across the three training and test conditions. Both pigeons and the model (when using lower-level DCNN representations) perform best when trained and tested on colored cardiograms. Training and testing on greyscale is intermediate. Training on color and testing on graysale is worst. The model, like the pigeons, shows a continuum of responses depending on the level of damage in the cardiogram (see Figure 4).

**Figure 9.**
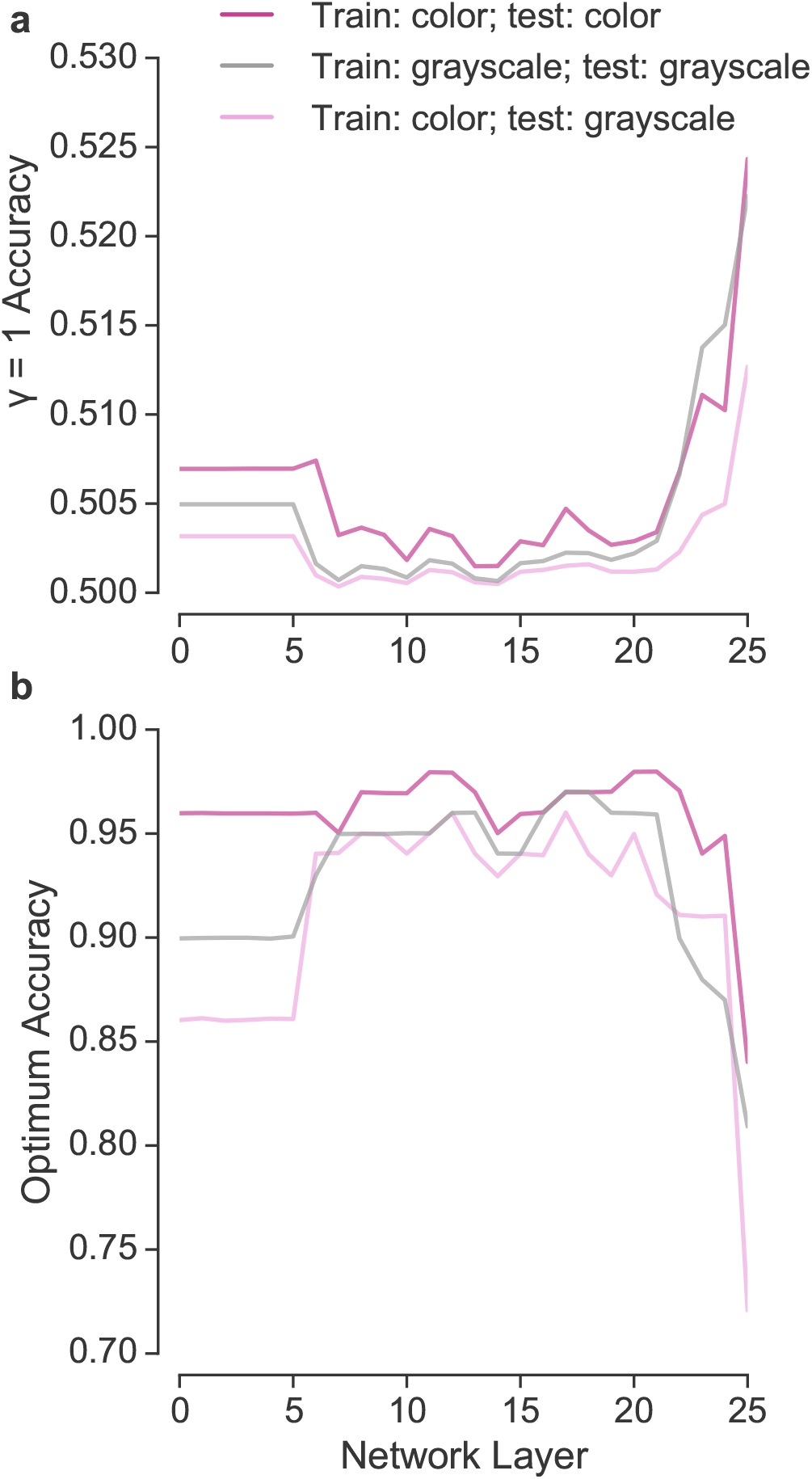
Accuracy of our model on each network layer when trained and tested in a manner analogous to the pigeons. The qualitative pattern of performance observed at the lowest network layers mirrors the performance of the pigeons. In panel **a**), *γ* is set to 1, whereas in panel **b**) an optimal boundary (determined from the training set) is used.

**Figure 10.**
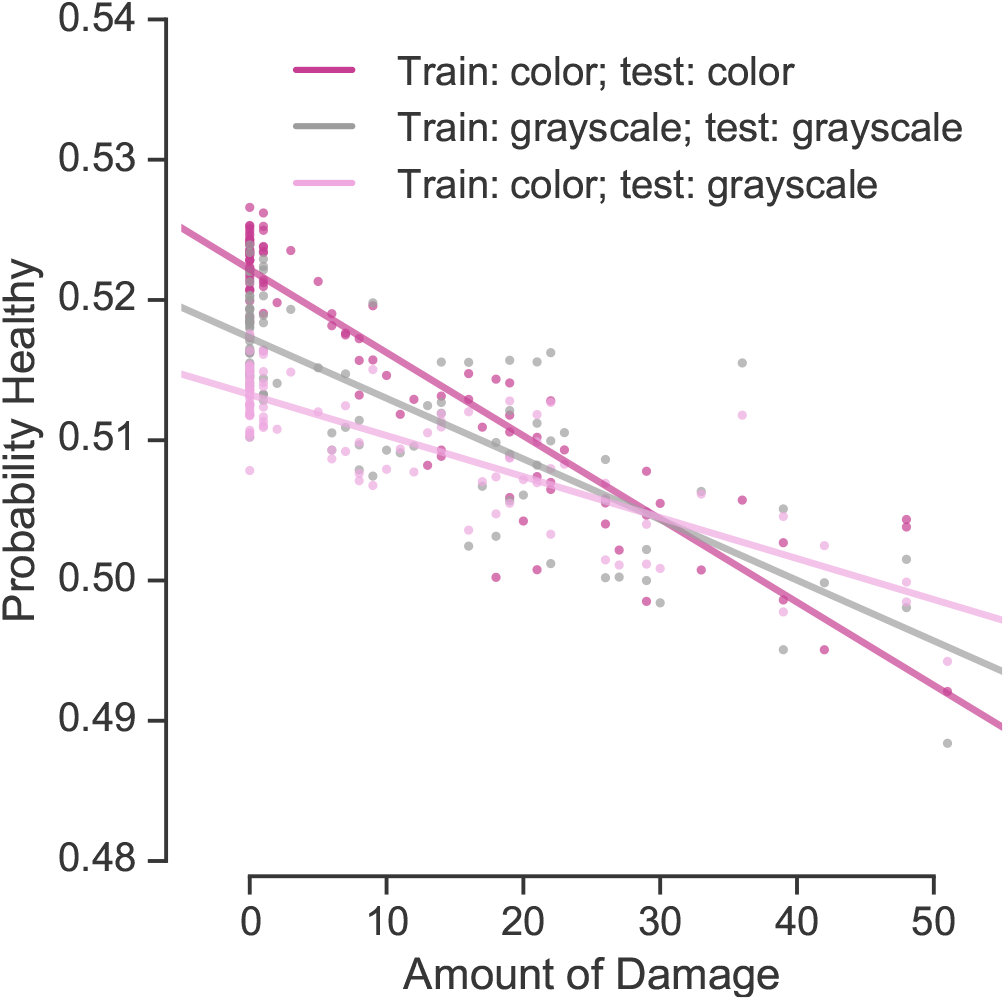
When relying on very low-level DCNN representations, the model, like the pigeons, shows a gradient of responding depending on the level of damage. As shown by the regression lines, the model displays the same difficulty ordering when generalizing to novel test stimuli. Training and testing on color stimuli leads to the best performance, whereas training on color stimuli and testing on grayscale stimuli leads to the worst performance.

The exemplar model using DCNN representations provides a tool to which to assess the nature of the representations used by learners, in this case pigeons. The modeling analysis suggests that the pigeons could be using very low (pixel-level) representations to support their performance. The modeling does not prove this conclusion, but does support it and demonstrates that such an account is plausible. The pigeon results diverge from our simulations of the shape bias in children (see Figure 2) which suggested that children rely on higher-level image statistics. Applying our model to such datasets provides insights not only into the populations considered, but into the types of knowledge required to perform well in various tasks.

## General Discussion

In this paper, we proposed an exemplar model of categorization that uses representations from a particular layer of a DCNN. Different model variants can be defined by relying on different DCNN layers. This approach has two primary strengths. First, by using representations from the DCNN, modelers do not need to rely on handcrafted and brittle representations. Instead, real-world stimuli, such as photographs, can be considered. Second, by comparing model variants, one can test hypotheses concerning the level of representation (i.e., which layer within the DCNN) a learner relied upon. Because the layers of DCNNs have been put in rough correspondence with processing stages of the ventral stream, these hypotheses extend to neural computation.

To make these model-based inferences stronger, we attempted to characterize the nature of the representations at various network layers. Our first simulation of the shape bias revealed that this bias relies on higher-level statistics. Indeed, when representations from lower layers of the DCNN were used, a color bias was observed.

A second set of simulations systematically evaluated the relative efficacy of various features at various network layers, as well as the importance of positional (i.e., location) overlap in images. From these simulations, we confirmed that shape becomes more prominent in network representations at more advanced layers, whereas stimulus dimensions like size and color relatively dominate at lower network layers. Many of these lower-level advantages are driven by positional overlap. At more advanced network layers, representations become increasingly invariant to location. Interestingly, we discovered that our model displays a preference for color over greyscale stimuli because of the similarity function used. Overall, these results are consistent with the idea that lower network layers are closely bound to the image whereas advanced layers can support more abstract representations.

In light of these results, one possibility is that higher-level features, such as shape, are represented at more advanced network layers whereas more basic features are represented lower in the hierarchy. An alternative possibility is that all features are more strongly manifested at more advanced network layers in absolute terms and that certain features, like shape, only come dominate in relative terms. In a third set of simulations with Gabor patches, we found support for the latter conclusion — although basic features first appear at lower network layers, they remain present in advanced network layers, as if the network operates as an inverted pyramid in which the range of represented features increases as one traverses layers.

With this understanding of network representations in hand and its relation to neuroscientific findings, we applied our model to a study in which pigeons learned to classify cardiograms as normal or abnormal. The goal was to determine whether pigeons relied on lower- or higher-level representations. We found that the pigeons performance was consistent with the use of very low-level representations that could be characterized as retinal. Our formal analysis parallels concerns of others with respect to pigeons’ (and other animals’) categorization limitations (e.g., Dittrich, Adam, Ünver, & Güntürkün, 2010; Lea et al., 2018; Linke, Bröker, Ramscar, & Baayen, 2017), despite what appears on the surface to be impressive feats of categorization. These results contrast with our shape bias simulations, which suggested human children rely on higher-level information. Interestingly, our model was able to predict pigeons’ preferences for colored cardiograms (in accord with human radiologists) based on the interaction between the multiple channel representation of color and the model’s similarity function. These simulations also serve to highlight the model’s utility in understanding what type of information is required to perform a task, which could be useful to experimentalists when aiming to design informative studies.

Although DCNNs can be treated as purely black-box solutions, our analyses suggest greater insight into behavior and the supporting neural representations can be gained by examining the bases for these networks’ performance (cf. MondragÃşn, Alonso, & Kokkola, 2017). Here, we focused on characterizing the information encoded at various network layers. We confirmed some intuitions, such as that shape relies on higher-level representations, whereas other conclusions were surprising. For example, we found support for the idea that purported lower-level features are retained at advanced network layers, a possibility that should be further pursued by neuroscientists (Ahlheim & Love, 2018; Hong et al., 2016; Lescroart & Gallant, 2019; Lu et al., 2018; Rice et al., 2014). The results presented here suggest a wealth of hypotheses that can be tested by comparing network representations to brain measures. Additionally, previous results in the literature aligning network representations and brain measures should be revisited in light of these new insights about network representations. Finally, our categorization model, which relies on DCNN representations, opens up a number of possibilities for learning studies using real-world stimuli.

## Notes

This work was supported by NIH (Grant 1P01HD080679), and a Wellcome Trust Investigator Award (Grant WT106931MA) to BCL, as well as The Alan Turing Institute under the EPSRC grant EP/N510129/1. Some of this work was originally reported at the 39th Annual Meeting of the Cognitive Science Society in 2017.

https://osf.io/jxavn/

